# Aerobic scope is sustained through a heatwave in juvenile Atlantic salmon

**DOI:** 10.1101/2025.07.03.662906

**Authors:** Lucy Cotgrove, Sergey Morozov, Miika Raitakivi, Evan Sala, Jenni M. Prokkola

## Abstract

Aquatic ectotherms are vulnerable to heatwave-induced physiological stress, which arises from increased energy demands and reduced dissolved oxygen content in warmer waters. Understanding thermal physiology is critical for predicting how commercially and ecologically important populations could be affected by the increasing risk of rising temperatures. Heatwave risk assessments often examine extremities of time scales: immediate impacts or long-term consequences. However, little is known about how consistently increasing mid-term thermal stress shapes aerobic performance in commercially important species such as Atlantic salmon (*Salmo salar*), which may face heat stress in rivers, especially at the juvenile life stage. By measuring how salmon juveniles manage their aerobic capacity at 16, 19 and 22°C using intermittent respirometry, we test if their thermal performance curve shows a decline at temperatures commonly occurring during heatwaves. Whole-animal metabolism was measured from control individuals kept at 16 °C before and after the heatwave, and after 4-5 days exposure at 19 and 22°C during the heatwave. We show that standard metabolic rate increases with temperature, but maximum metabolic rate and aerobic scope do not change between these temperatures. These findings suggest that juvenile Atlantic salmon may have limited capacity to increase aerobic performance during moderate heatwaves, leaving them vulnerable to cumulative effects of oxygen limitation to vital functions such as growth and stress responses. As climate change intensifies, incorporating thermal performance curves into conservation strategies can be utilized for predicting population resilience and informing effective management.

**Lay Summary:** In juvenile Atlantic salmon, standard metabolic rate increases but there is no difference in maximum metabolic rate and aerobic scope with increasing temperatures. Considering the increased metabolic costs of activities at higher temperatures, our results show that juvenile salmon are vulnerable to deteriorating performance at temperatures commonly experienced in present-day heatwaves.

## Introduction

With climate change increasing the severity and frequency of heatwaves (IPCC, 2023), organisms may experience physiological thermal stress and even mortality due to extreme temperature fluctuations. Understanding organisms’ thermal physiology is therefore critically important for conservation (Clark et al., 2013; Deutsch et al., 2008). Heatwaves frequently surpass the temperature range that species have been adapted to, often cumulatively and repeatedly, imposing selection on thermal performance. This is especially true for ectotherms, whose internal temperature are regulated by the surrounding environment. Some species may demonstrate adaptive or resilient responses to heatwaves including behavioural adjustments, physiological acclimation, or shifts in geographical ranges (Breedveld et al., 2023; Burraco et al., 2020; Jørgensen et al., 2022). However, these responses may not always be sufficient to counteract the negative consequences of extreme weather fluctuation (Perry et al., 2005; Pörtner & Farrell, 2008). Most physiological processes in ectotherms, including development, reproduction, and metabolism, are temperature-dependent (Angilletta et al., 2010; Hochachka & Somero, 1973; Schulte, 2015; Seebacher, 2009). These processes can be described using thermal performance curves (TPCs), which illustrate how a biological rate, such as oxygen consumption or activity per unit time, changes with temperature (Schulte et al., 2011).

TPCs can be estimated using thermal windows, critical temperatures, and preferred temperature ranges (Jørgensen et al., 2021; Ørsted et al., 2022; Peralta-Maraver & Rezende, 2021). TPCs can be used to compare inter- and intraspecies responses to anticipated temperatures under a future climate and how this might vary across life stages (Lefevre et al., 2021; Seebacher & Little, 2021). In water-breathing aquatic ectotherms, such as fishes, TPCs linked to demand, uptake and delivery of oxygen are particularly important because of the decreasing solubility but increasing oxygen consumption in warmer water (Little et al., 2020; Little & Seebacher, 2021; Seebacher & Little, 2021; Verberk et al., 2022). Consequently, TPCs related to metabolic rates and aerobic performance are pivotal to understanding the effects of heatwaves on fishes.

Metabolic rate is a fundamental aspect of fish physiology, reflecting the energy expenditure associated with basic life processes such as respiration, digestion, and movement (Brett, 1964; Fry et al., 1947). Standard metabolic rate (SMR) of an animal represents the baseline rate of aerobic metabolism required to sustain life in a post-absortive, resting state, while the maximum capacity for aerobic performance is set by the maximum metabolic rate (MMR) (Sandblom et al., 2016). The difference between SMR and MMR represents the aerobic scope (AS) of an animal and theoretically determines the capacity of aerobic metabolism to support key life-history attributes such as activity, growth and reproduction, each of which has a specific oxygen cost (Fry, 1971; Fry et al., 1947). Therefore, reductions in aerobic scope can impair physical performance (Johansen & Jones, 2011; Pörtner & Farrell, 2008; Priede, 1985). Given that the SMR of ectotherms rises with temperature and often no such increase is detected in MMR (Fry et al., 1947; Norin et al., 2014; Sandblom et al., 2016), increases in temperature beyond optimum can result in a decline in AS (Farrell, 2016; Fry, 1971; Lefevre et al., 2021). However, maintaining growth and other aerobic functions with increasing temperature requires an increasing AS, because of the higher costs of activities such as digestion and assimilation and swimming (Jahn & Seebacher, 2022; Jutfelt et al., 2021). When AS declines at increased temperatures, this may cause declining growth, health, or survival of individuals (Alfonso et al., 2021; Jutfelt et al., 2021; Navarro et al., 2019; Sadoul & Vijayan, 2016). Alternatively, AS can plateau at higher temperatures suggesting that oxygen supply can limit performance as temperature increases (hvas et al 2017, Závorka et al., 2020, Christensen et al 2012). As AS plateaus but SMR and the costs of aerobic activities continue to increase, fish might become more constrained in their physiological state and experience cumulative thermal stress, leading to mortality. Both declines and plateaus of AS pose significant risks for salmonids, which include economically important, cold-water adapted species. In these fish, limitations of AS has already been linked to reduced growth, survival, or collapse of aerobic capability during heatwaves (Hvas 2017 (Eliason, Wilson, et al., 2013; Pörtner & Farrell, 2008; Wade et al., 2019).

The Atlantic salmon (*Salmo salar* L.) is faced with supra-optimal temperatures especially at riverine life stages, including juveniles (parr) and adults during their spawning migration, with reports of present-day river temperatures up to 28 °C (O’Sullivan et al., 2023; Strøm et al., 2019). Juvenile Atlantic salmon rear for one to five years in freshwater before migrating to the sea (Aas et al., 2010), with potential to encounter multiple heatwaves. These can have direct effects on survival, but also cascading effects on population dynamics through reduced growth rates. Faster freshwater growth in salmon is linked to higher survival and faster maturation at sea (Hutchings & Jones, 1998; Simpson, 1992; Thorpe, 2007). In some populations, a negative association between freshwater growth and river temperature has been detected (Alioravainen et al., 2023). For the majority of their life cycle at sea, Atlantic salmon will experience temperatures below 8°C (Jensen et al., 2014; Lacroix, 2013; Reddin, 1985). Atlantic salmon tend to avoid temperatures above 15°C (Fisher & Elson, 1950; Johansson et al., 2009; Lacroix, 2013), and their optimal temperature range for feeding and normal behaviour in the wild has been suggested to be 6–20°C, with peak growth rates occurring at 16–17°C in aquaculture conditions (Dwyer & Piper, 1987; Elliott, 1982). Previous work assessing thermal performance in Atlantic salmon has often focussed on upper limits during acute exposure of CTmax experiments under less than 24 hr(Desforges et al., 2023 and papers within), or long-term stable temperature increases over several months (Anttila et al., 2014; Del Rio et al., 2021; Hvas et al., 2017). When aerobic scope has been measured in a heatwave-context in a related species, (Casselman et al., 2012) found AS peaked at 17 °C, and decreased until testing finished at 21 °C in juvenile Coho salmon *(Oncorhynchus kisutch*). In post-smolt Atlantic salmon, AS tended to increase (a non-significant effect) as temperature increased from 13 °C to 23 °C, but swimming ability and feeding rate were greatly decreased and mortality increased, when metabolic rates of salmon were tested in groups without a cumulative exposure (Hvas et al., 2017). Further, thermal variability in acclimation regime had no effect on salmon AS compared to a stable acclimation when metabolic rates were tested acutely at the same temperature (Morissette et al., 2021). Still, information on the cumulative effects of thermal fluctuations over several days or weeks, which better reflect natural heatwave patterns, are limited (Morash et al., 2021; Nuic et al., 2024). This information is pivotal for the conservation of Atlantic salmon. The diminishing of salmon populations has already reduced opportunities for commercial and recreational fishery, cultural traditions, and increased the risk of local extinctions (ICES, 2024, Dadswell et al., 2022)

Here, we test how mid-term increasing temperatures that are prevalent in rivers during heatwaves impact metabolic rates and AS in hatchery-reared juvenile Atlantic salmon. We hypothesise that SMR will increase as temperature rises. We expect MMR to show little variation between temperatures, and therefore a decrease in AS as temperature increases. These findings will provide insight into the physiological constraints juvenile salmon may face during heatwaves, using an ecologically relevant temperature regime.

## Methods

### Fish Husbandry

The experiment was performed with a permit granted by the Finnish Project Authorisation Board (no. ESAVI/16748/2023). Families of Atlantic salmon were bred using a hatchery broodstock originating from River Iijoki, maintained by Natural Resources Institute of Finland (Luke), in Taivalkoski, Finland. The broodstock was established in 2012 from parents that were originally stocked into the river as juveniles/smolts, and therefore had completed half of their life cycle in the wild. Individuals from the broodstock were crossed to make full-sib families of offspring in October 2022. The embryos were incubated in Taivalkoski hatchery in family-specific flow-through trays at natural temperatures of incoming water from River Ohtaoja, and dead embryos were removed regularly.

Embryos were disinfected by iodine bath (Buffodine) on 9th March 2023, and transported to a LUKE hatchery in Laukaa, Finland, where experiments were conducted. Survival between transportation and start of the experiment was 22%. Excess mortality may have been caused by the closeness to the embryo’s hatching date and transporting to the fish farm (no disease was observed in the offspring). Embryos from three unrelated families were incubated and reared in separate circular tanks (diameter 80 cm) and supplied with a continuous flow of filtered water from a mix of local lake Peurunkajärvi and river Peurunkajoki. Feeding of alevins was started with powdered feed (BioMar Group) when most of the egg yolk had been consumed at approximately 25 May, 2023. Tanks were cleaned by scrubbing surfaces and siphoning excess food twice a week until 10 June, 2023, and then daily until the end of the experiment. Feed rations were calculated from growth predictions assuming feed conversion efficiency of 0.8 (Elliott & Hurley, 1997), and adjusted such that a small amount of feed was left uneaten. Fish were fed using 12h belt feeders with the daily ration, except for one day preceding metabolic measurements, when fish were fasted. During the summer and the experiment, fish were reared under constant light. Rearing temperature raised gradually with natural water temperatures over approximately three months until the experiment was started; temperature varied from 9.2 to 15.7 °C between first feeding and experimental period.

At the start of the experiment 27 July, 2023, families were divided into two groups, control and heatwave, with approximately same number of individuals in each group within a family (N = 24 to 62 within family/tank). Fish were kept in three tanks each for control and heatwave groups. Densities within families were kept similar between control and heatwave tanks throughout the experiment by simultaneously removing fish from control tanks to match densities in of heatwave tanks. Control tanks were kept at close to 16 °C (mean = 16.1, SD = 0.5) throughout the experimental period by regulating the incoming water flows from the river and the lake with a natural temperature difference. Temperatures in one control tank (the same water source used in all three tanks) and the three heatwave tanks were measured every 30 min with Hobo 64K Pendant temperature loggers (Hobo, Bourne, MA, USA).

Temperatures of the heatwave tanks were regulated via a heating tank to offset the colder temperatures of the room and incoming water using a temperature controller (TS1000, H-Tronic GmbH, Germany) with a temperature probe PT-1000 located in each tank, and an Eheim 600 or 300 pump (Eheim, Deizisau, Germany) depending on the distance from the heating tank. The pumps circulated water through stainless steel coils in the heating tank, in which temperature fluctuated between 30-35 °C. Heatwave tank water was aerated constantly to prevent declines in dissolved O_2_ related to increasing temperatures, as the flow-through rate of fresh water to the tanks was reduced (approx. 1 L min^−1^) during the heatwave to maintain temperatures.

### Experimental Heatwave

Fish in the heatwave treatment were exposed to three test temperatures: 16 °C, 19 °C, and 22 °C (Figure 1). First, 16 °C was maintained as a control temperature for eight days, after which 3-8 individuals from each family were randomly picked for measurements of metabolic rate (similar sample size across temperatures within a family), and moved into a fasting tank, where they were unfed until the measurements started the following day. Temperature of the fasting tank was the same as the heatwave tanks. After the measurement of MMR and SMR at 16 °C, temperature was increased by one degree per day for three days, and maintained for 4-5 days, after which metabolic rate measurements were taken from a new set of fish at 19 °C. This was repeated and metabolic rate measurements were taken at 22°C. Due to a malfunctioning temperature controller at 19 °C (after moving fish to acclimation for B3 and B4), there was an increase in temperature in one of the heatwave tanks to 21.5 °C (Figure 1). Metabolic rate experiments were conducted once per individual, and fish were euthanised after the SMR measurement.

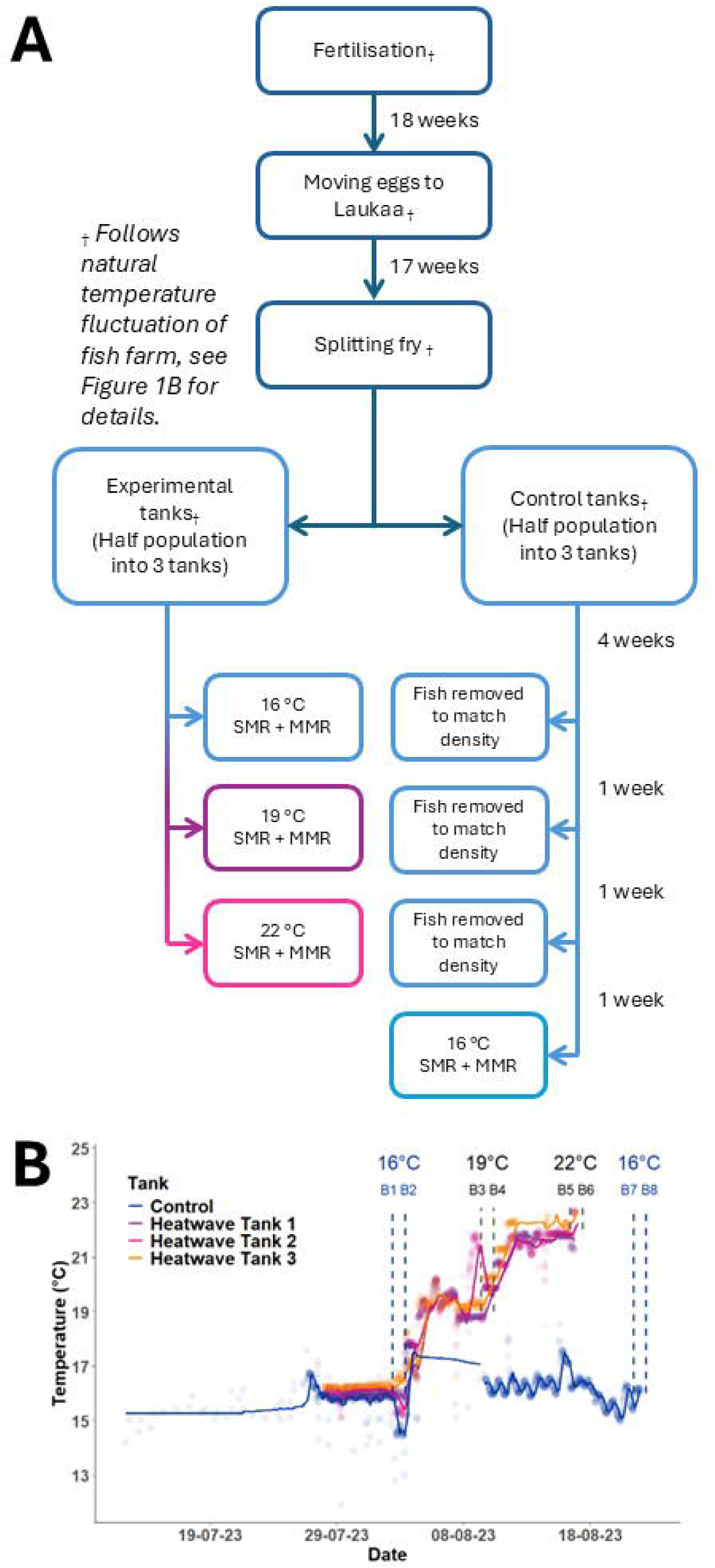

To measure MMR, fish were transferred from the fasting tank into 50 L buckets filled with 5 cm of water at the testing temperature using plastic cups of water to reduce air exposure. Fish were encouraged to swim by hand around the circular container (Prokkola et al., 2022; Raby et al., 2020). Chasing lasted 2 min, and all fish were unresponsive to the touch of caudal fins, thus determined fatigued. The fish were then rapidly transferred to respirometry chambers using plastic cups with water. Two people chased one fish each at once, and each batch of max. 16 individuals was processed within 2 h. Fish were left in respirometry chambers overnight in order to capture the MMR and SMR for each fish, as per (Chabot, McKenzie, et al., 2016). Chambers were flushed with fresh water from the tank for 5 min every 15 min, allowing fully aerated water to enter the chamber. The lowest recorded oxygen level in any chamber was 6.7 mg L ^−1^ at 22 °C. After SMR measurements, fish were euthanised using an overdose of MS-222, measured, and weighed.

Immediately before and after metabolic measurements, background respiration was measured in empty chambers for three measurement cycles (5 min flush, 15 min measurement) and an average of these slopes was used to adjust the data for MMR and SMR. For those trials which took place immediately after the previous, the post-trial background data of the previous trial was used as the pre-trial background values. To prevent bacterial build-up, the system was bleached between temperature changes, and all chamber parts scrubbed to limit bacterial growth and rinsed thoroughly. Additionally, the water in the respirometry tank was continuously circulated through a UV filter.

A total of 120 fish were subjected to intermittent flow respirometry after a 24h fasting period to provide estimates of metabolic rate (SMR, MMR, AS see table S1) (Killen et al., 2021; Svendsen et al., 2016). Due to death in respirometer chambers, exclusion due to outliers, incomplete data recording, and due to computer failure, 72 fish were included in the analysis (Table S2).

### Respirometer design

For a summary of the respirometry design and measurement protocol, see Supplementary Table 1, after (Killen et al., 2021). A 16-chamber intermittent flow respirometer, each chamber containing an individual fish, was submerged in a temperature-regulated water bath (16, 19 & 22 ± 0.1 °C; 200 L) that was saturated with air. The oxygen content of the water in the chambers was measured every two seconds using a 4-channel fibre optic oxygen meter with associated oxygen sensors and software (FireStingO2; PyroScience GmbH, Aachen, Germany). All 16 O_2_ sensors were calibrated for 0% oxygen saturation simultaneously with O_2_-free water, made using sodium sulphate, and simultaneous for 100% oxygen saturation with air-saturated water using an air stone before the first measurements at each temperature. Temperature-compensation for O_2_ saturation was based on PT100 temperature sensors, which were placed in the middle of the respirometer tank and connected to each of the Firesting meters. The chambers were shielded from disturbance and light using an opaque plastic cover during the measurement of SMR. Flush and recirculation pumps were controlled using PumpResp controllers (4-channel model, FishResp, Finland, https://github.com/embedded-sergey/PumpResp, (2024)). The temperature of the respirometer tank was maintained using a temperature controller (as in the heatwave tanks) and a reservoir connected to a TECO 2000 Chiller/heater. The reservoir received constant inflow of water (approx. 1 L min^−1^) from the same source as the rearing tanks to maintain good water quality. Chambers for the respirometer were made using glass tubing (120 mm length, inner diameter 38 mm, wall thickness 3.2 mm) and plastic caps that were 3D-printed on PA2200 polyamide. The ‘HeiBer’-caps were designed by Heidrikur Bergsson, University of Copenhagen (https://zenodo.org/record/4062429#.YMSW7h1RVTY). Volume of chambers and tubing was 131.86 ± 1 mL (the exact volumes per chamber are reported in an online data repository, see Data Availability). Water inside the chambers was mixed by a plastic disk attached to each cap to distribute the flow. The disks were 3D printed using the same method as the respirometry caps and attached with stainless steel screws. Caps were sealed using rubber O-rings and connected to the flush and recirculation pumps and valves with gas-impermeable Tygon tubing (Tygon S3 E-3603, Satint-Gobain, Paris. France). The recirculation system was confirmed to be waterproof by filling with water and plugging the flush pump and probe connections. Water was recirculated using one submersible pump (12V DC, 6W 2 L min^−1^ pump, Qingdao Xinhui Hardware Machinery Co., Ltd., Qingdao) per channel within the recirculation loop. Chambers were flushed using a second pump (same as previous), using an inflow of water from the tank into the chambers, and excess water was flushed through the chamber to a flush pipe which was placed with the end above the water surface. This allowed fully aerated water to enter the chamber. Flow rate through both the flush line and recirculation loop could be controlled by the valves attached within the flow. Flow rate of the respirometers was approximately 0.1 (± 0.008) L min^−1^ without fish. Information about time and pump phase (either flush or measurement) was recorded by the PumpResp controllers and stored on a computer. Flow had no apparent effect on movement of fish within the chambers in SMR or MMR measurements.

### Data Analysis

All data and statistical analyses were done in R environment (R Core Team, 2022). The FishResp package (Morozov et al., 2019) was used to filter and calculate metabolic rate estimates. Slopes of oxygen consumption were adjusted for bacterial oxygen consumption using pre-and post-background measurements, assuming a linear change. For SMR, oxygen consumption (mg O_2_ h^−1^) for each measurement phase was derived from the slope of linear regression of dissolved oxygen concentration over time. The SMR slopes were quality filtered and smoothed before analysis to exclude non-linear declines of oxygen: first, slopes were filtered based on R^2^ > 0.95, then slopes that did not meet the R^2^ criteria were smoothed using a running mean of 29 seconds (Chabot et al., 2021), then R^2^ filter was re-applied, and finally, linearity was checked visually from all slopes that met the criteria of R^2^ > 0.95. The first and last 60 seconds of measurement periods were excluded to account for mixing of water and changes in flow due to flushing. The mean of the lowest normal distribution (MLND) was used to estimate SMR for each individual from the extracted slopes (Chabot, Steffensen, et al., 2016). Slopes for MMR were extracted using a derivative of a polynomial curve fitted on each measurement (function *smooth.spline,* d.f. = 10) as in the *spline-MMR* method (Prokkola et al., 2022), i.e. from the beginning of the MMR measurement, when respiration was the highest. The slopes were then used to calculate MMR (mg O_2_ h^−1^) using *FishResp*. AS was calculated as the difference between the absolute MMR and SMR.

A linear mixed effect model was fitted (estimated using restricted maximum likelihood and *nloptwrap* optimizer) to predict SMR, MMR and a non-linear mixed effect model was fitted to predict AS, all of which were log transformed. Temperature and log-transformed mass as fixed effects (formula: Metabolic Measurement ∼ Temperature + Mass). The model included batch as a random effect (formula:∼1|Batch) (Zuur et al., 2009). Homoscedasticity and normality of residuals were assessed by visual inspection of residual plots. 95% Confidence Intervals (CIs) and p-values were computed using a Wald t-distribution approximation. Post-hoc comparisons of temperatures were performed using Tukey method of comparing estimates, and P value < 0.05 was considered significant. Statistical models were fitted using *nlme*, *lme4* and *lmerTest* packages in R, and post-hoc analysis was performed in *emmeans* package (Bates et al., 2015; Kuznetsova et al., 2017; Lenth, 2017; Pinheiro et al., 1999). For visual representation, SMR, MMR, and AS data were mass-adjusted to account for the hypo-allometric scaling of metabolic rates by linear regression of log_10_-transformed metabolic rates against log_10_-transformed body mass (Figure 2). Finally, Pearson’s correlations were calculated among the either absolute or mass-adjusted metabolic variables.

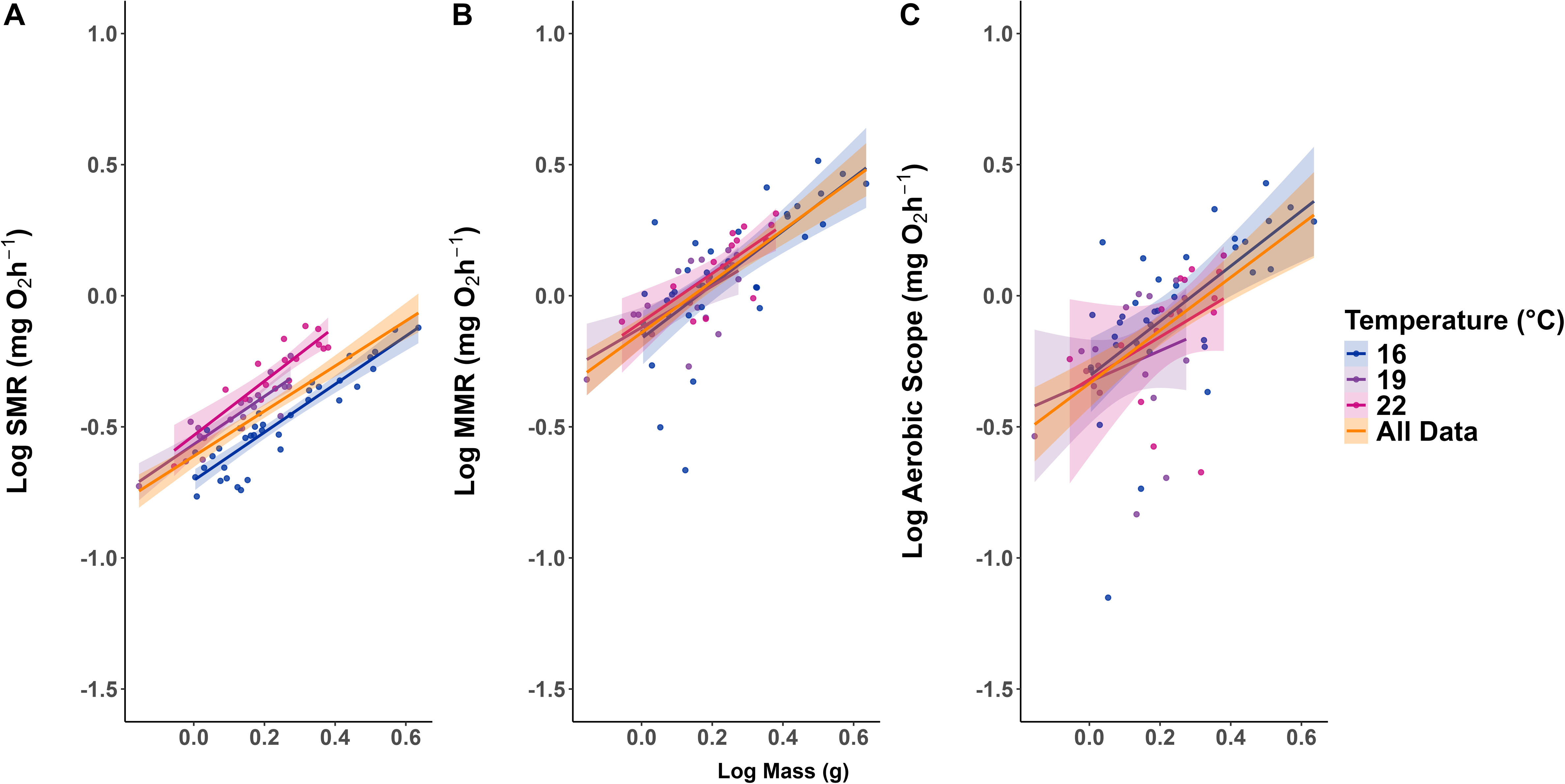

## Results

In total, the data of 72 fish was used at three temperatures across four consecutive weeks: 16°C, 19°C, and 22°C and 16 °C again (Table 1, Table S3). The metabolic rates and AS scaled positively with body mass, with scaling exponents from 0.86 to 1.1 (Figure 2, Table 2, Table S4).

**Table 1:** Descriptive statistics for Mass (g) and Length (mm) for juvenile salmon tested at different temperatures. n = number of individuals, xl = mean metabolic rate, σ = standard deviation, Min = minimum value, Max = maximum value.

| *Temperature (°C)* | *n* | *Mass (g)* |  |  |  | *Length (mm)* |  |  |  |
| --- | --- | --- | --- | --- | --- | --- | --- | --- | --- |
| | | $\bar{x}$ | $\sigma$ | *Min* | *Max* | $\bar{x}$ | $\sigma$ | *Min* | *Max* |
| 16 (week 1) | 22 | 1.42 | 0.41 | 1.01 | 2.59 | 57.64 | 5.38 | 50 | 69 |
| 19 | 19 | 1.35 | 0.35 | 0.70 | 1.88 | 56.84 | 4.71 | 47 | 64 |
| 22 | 16 | 1.79 | 0.42 | 0.88 | 2.40 | 61.00 | 4.35 | 54 | 68 |
| 16 (week 4) | 15 | 2.56 | 0.88 | 1.40 | 4.32 | 68.53 | 8.22 | 56 | 82 |
| *Total* | 72 | 1.72 | 0.70 | 0.70 | 4.32 | 60.44 | 7.16 | 47 | 82 |

**Table 2:** Model summaries for metabolic rate estimates, including fish body mass and temperature (16, 19 and 22 °C) as independent variables. Batch is included as random effect. Brackets indicate standard error, asterisks indicates significance values with *p<0.05; **p<0.01; ***p<0.001).

|  | *log10(SMR)* |  |  | *log10(MMR)* |  |  | *log10(AS)* |  |  |
| --- | --- | --- | --- | --- | --- | --- | --- | --- | --- |
|  | *Est.* | *CI* | *p* | *Est.* | *CI* | *p* | *Est.* | *CI* | *p* |
| log10(mass) | 0.922 | 0.057 | <0.001 | 0.978 | 0.120 | <0.001 | 1.044 | 0.229 | <0.001 |
| 19 °C | 0.131 | 0.027 | 0.004 | 0.011 | 0.046 | 0.826 | -0.038 | 0.086 | 0.674 |
| 22 °C | 0.198 | 0.027 | <0.001 | 0.040 | 0.046 | 0.424 | -0.033 | 0.086 | 0.717 |
| Intercept | -0.704 | 0.021 | <0.001 | -0.152 | 0.038 | 0.004 | -0.342 | 0.072 | <0.001 |
| ***Random Effects*** |  |  |  |  |  |  |  |  |  |
| Intercept SD (Batch) |  | 0.0215 |  |  | 0.0116 |  |  | 0.0116 |  |
| Residual SD |  | 0.0638 |  |  | 0.1492 |  |  | 0.1492 |  |
| Marginal R<sup>2</sup> |  | 0.849 |  |  | 0.521 |  |  | 0.521 |  |
| Conditional R<sup>2</sup> |  | 0.864 |  |  | 0.524 |  |  | 0.524 |  |

We found temperature to be a significant predictor of SMR (16–19 °C: t(66) = 4.80, P < .001; 16–22 °C: t(66) = 7.43, P < .001, Table 3). There were significant differences in SMR between all temperature groups, except between 19 °C and 22 °C (Figure 3, Table 4). There was a 57% increase in predicted SMR for 6 °C increase in temperature (Table 4). However, there was no significant change in MMR or AS between temperatures (Figure 3, Table 2).

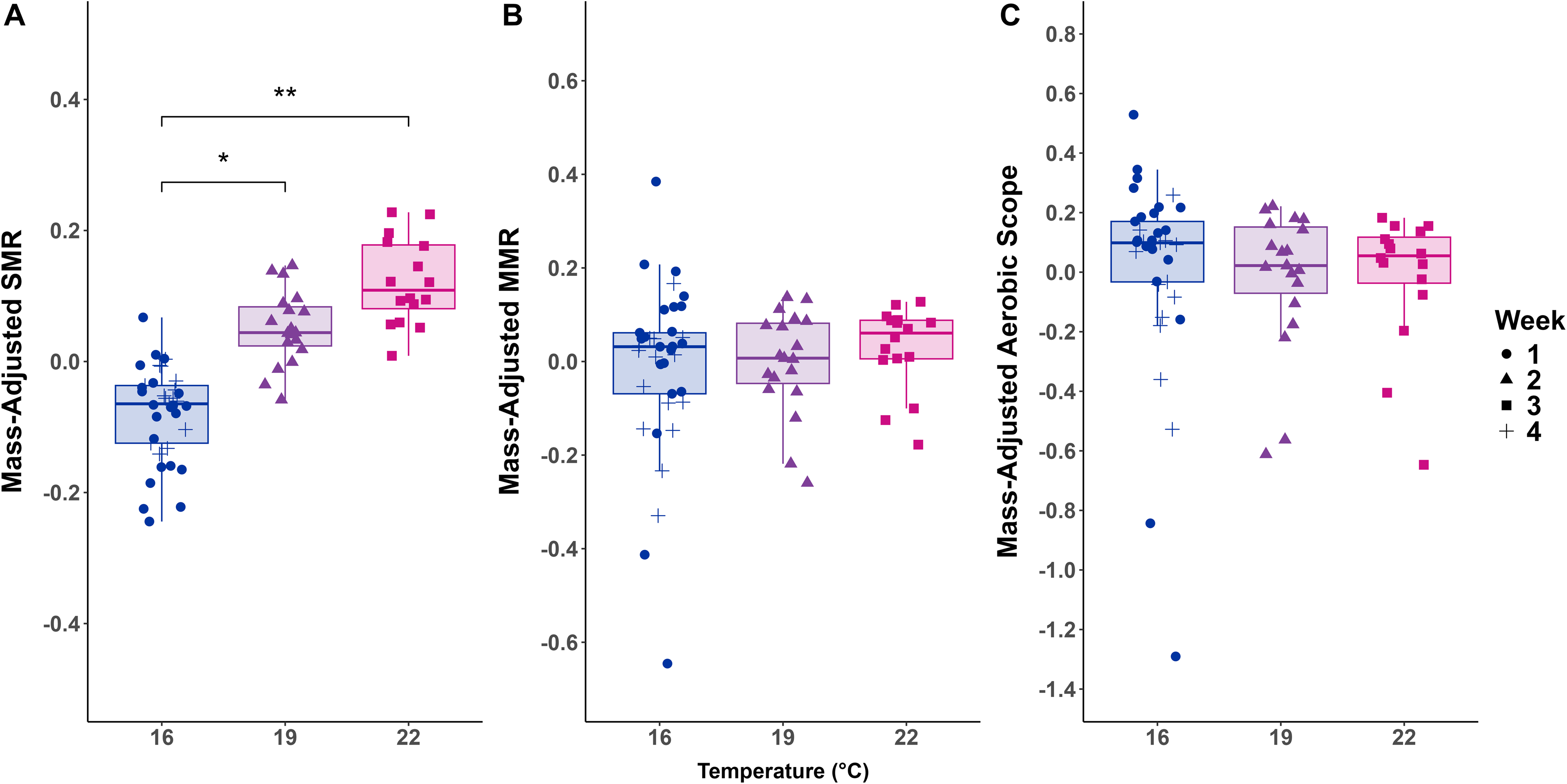

**Table 3:** Post-hoc ANOVA contrasts from the results of the SMR model described in Table 2, comparing the temperature treatments 16 °C, 19 °C, and 22 °C. SE shows standard error, df degrees of freedom and the associated *t* ratio and *p* (significance) with *p<0.05; **p<0.01; ***p<0.001.

| *Comparison* | *Estimate* | *SE* | *df* | *t* | *p* |
| --- | --- | --- | --- | --- | --- |
| 16 °C - 19 °C | -0.13 | 0.03 | 4.66 | -4.71 | **0.014** |
| 16 °C - 22 °C | -0.20 | 0.03 | 4.73 | -7.42 | **0.002** |
| 19 °C - 22 °C | -0.07 | 0.03 | 4.94 | -2.13 | 0.178 |

**Table 4:**
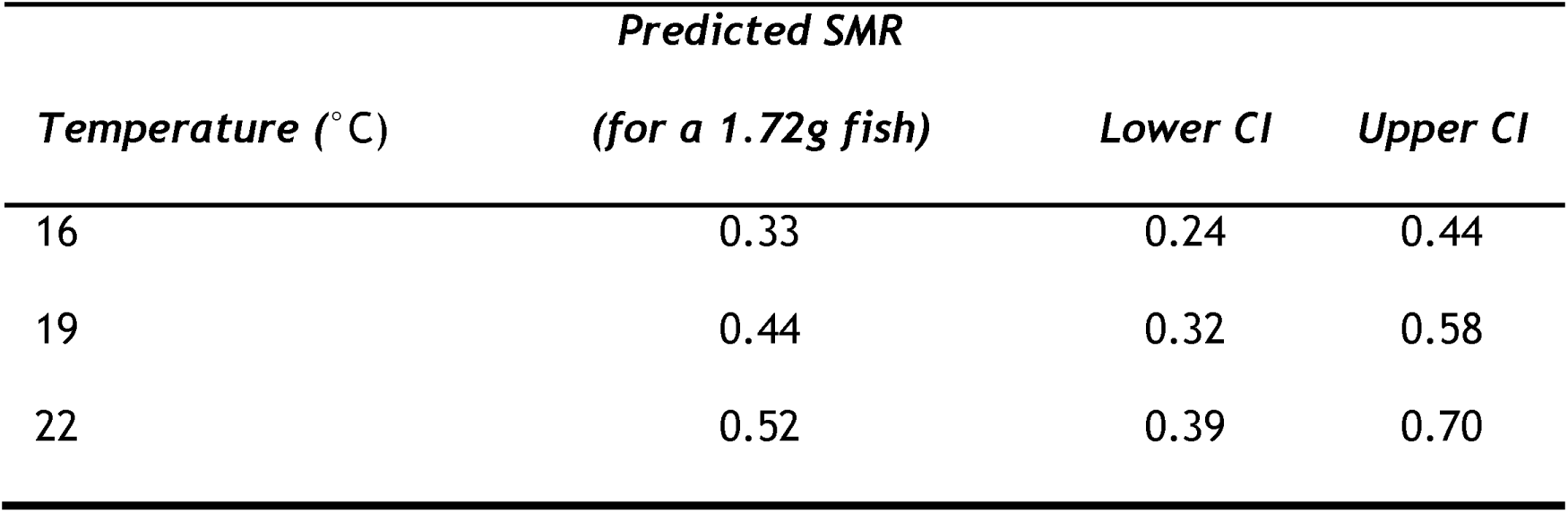
Table showing predicted SMR (mg O_2_ h^−1^) and 95% confidence intervals (CI) for a mean mass fish (1.72g) at different temperatures.

### Correlations among metabolic variables

At all temperatures, MMR and AS were highly positively correlated (r > 0.9) in both raw and mass adjusted data. Absolute SMR and MMR were positively correlated at 16 and 22 °C. Absolute SMR and AS were positively correlated at 16 °C. However, mass adjusted SMR and AS were negatively correlated at 19 and 22 °C (Table S5, Figure S1).

## Discussion

This study demonstrates that rising temperatures lead to an increase in standard metabolic rate (SMR) in juvenile Atlantic salmon. However, there was no change in maximum metabolic rate (MMR) or in aerobic scope (AS) across the tested temperatures. While an increase in SMR with temperature is well established in ectotherms due to the effect of temperature on biochemical rates (Clark et al., 2013; McKenzie et al., 2021; Raby et al., 2016), there are more mixed results on the relationship of MMR, AS and increasing temperatures in fishes.

The TPC of AS in fish has been suggested to be a bell shaped curve (Pörtner & Farrell, 2008). For example, in pink salmon (*O. gorbuscha*), AS peaked at 21 °C and declined with further warming, although this temperature increase occurred over a maximum of 2 days (Clark et al., 2011). Other studies note a decrease in AS, which could elude to the post peak decrease of the AS curve, as these are tested at higher temperatures or after longer acclimations, for example, in a recent study on salmon parr, AS decreased with temperature from 15 to 21°C after a three-week acclimation (Nuic et al., 2024). This could suggest long term exposure to a heatwave leading to aerobic collapse. In contrast in murray cod (*Maccullochella peelii peelii*) and temperatures that increased from 14 – 29 °C, where fish were acclimated for at least 18 h to each increase, resulted in a continuous increase in aerobic scope (Clark et al., 2005).

Other studies report a plateau of AS at high temperatures. For example, there were no differences in AS between acclimation temperatures in minnows (*Phoxinus phoxinus*) after eight months (Závorka et al., 2020). Morissette et al. (2021) found that below 23 °C, there was no difference between any metabolic rates after different temperature acclimations, however in their study all metabolic rates were measured at the same temperature, which complicates the interpretation of the results. In gobies acclimated for three weeks, AS was seen to plateau between 15 and 28 °C after initially increasing (Christensen et al., 2021). Further, in Atlantic salmon post-smolts, aerobic scope was increased with 4-week acclimations at temperatures from 3– 13 °C, and plateaued from 13 °C up to a temperature of 23 °C. Interestingly, fish condition and swim speed dropped and mortality increased (Hvas et al., 2017). Upper thermal limit of salmon has been widely reported at 23 °C (Elliott & Elliott, 2010), and notable, often cited work (Jensen et al., 1989) found optimal salmon temperature for growth to be 16 °C. Our results agree with Hvas et al. (2017) as we found that AS plateaued at 16–22°C when measured at acclimation temperature, and it is likely that oxygen supply can limit performance of salmon as temperature increases. An absence of increase suggests physiological systems supporting oxygen uptake such as cardiac function, gill function and haemoglobin oxygen affinity are no longer able to scale with increasing thermal pressure (Anttila et al., 2014; Schulte & Healy, 2022). As AS plateaus but SMR and the costs of aerobic activities continue to increase, fish might become more constrained in their physiological state and experience cumulative thermal stress, leading to mortality.

AS is proposed to be of ecological significance because it defines the upper limit for oxygen allocation by a fish to sustain aerobic activities such as foraging, digestion, tissue deposition, migration, reproduction, and so forth (Claireaux & Lefrançois, 2007; Farrell et al., 2009b; Fry, 1971; H.-O. Pörtner, 2010; Schulte, 2015). For example, reduced appetite could be behaviourally driven to avoid anaerobic metabolism, as suggested by the reduction of feed intake during low AS (review by (Jutfelt, 2020). In support of this, reduced growth and lower condition factor at higher temperatures has previously been documented in Atlantic salmon, when comparing increases to temperatures up to 19 °C (Hevrøy et al., 2015; Kullgren et al., 2013). By measuring AS as an indicator of thermal performance, we can predict that at high temperatures, Atlantic salmon in the wild would either have to seek thermal refuge, or compromise aspects of their performance in order to cope with the high temperatures, which could result in poor growth or health (Koskela et al., 1997).

It is important to note that both TPCs and upper thermal limits are dependent on acclimation duration and rate of warming (Currie et al., 2014; Lefevre et al., 2021; McKenzie et al., 2021). Most research in fishes has been conducted with stable thermal profiles, whereby each individual experiences only one acclimation temperature (Hvas et al., 2017). In contrast, our study accounts for a cumulative effect of exposure to higher temperatures, using a gradual heating regime. While some acclimation is expected in the MMR over short-term temperature increases ((Norin & Clark, 2016); and papers within), each temperature exposure in this study occurred over 10 days, which may not be enough time to see notable plasticity in salmon’s metabolic ceiling. Sandblom et al. (2017) found no difference in MMR or AS between acute (24 h) or chronic warming (years) regime in perch (*Perca fluviatilis*), but that all warm-acclimated fish (5 days acclimation, 23 °C) have a higher MMR than fish acclimated at 18 °C, their reference temperature. However, an ecologically relevant scenario for salmon in their environments is a heatwave lasting from a few days to few weeks, thus these time frames are the most relevant to study from a conservation perspective. By testing the cumulative effect of a heatwave mirroring the temperatures seen in the wild, we can better predict how salmon could cope in a realistic exposure to sub-lethal temperatures (Morash et al., 2018).

The general expectation is that longer acclimation with a slower increase of temperature should be beneficial to survival, giving the fish time to make compensatory physiological modifications such as displaying plasticity in SMR or MMR, enabling AS to be maintained over a broad range of temperatures. However, in barramundi MMR and AS increased when exposed to acute 10 °C warming, due to a higher increase in MMR than SMR, but after 5 weeks of acclimation, AS was similar, mainly due to a reduction in MMR (Norin et al., 2014). A very similar pattern was observed in black sea bass (*Centropristis striata)* and common triplefin (*Forsterygion lapillum*), where short term exposure caused a raise in AS, before it decreased after some weeks of exposure (Khan et al., 2014; McArley et al., 2017; Slesinger et al., 2019). Additionally, in salmon post smolts, post-stress peak cortisol levels significantly increased with higher temperatures (Madaro et al., 2018). This suggests that despite a stable aerobic capacity to cope with temperature increases, costly physiological mechanisms may be occurring, and therefore a plateau in AS does not necessarily suggest a broad optimum temperature range.

MMR did not differ across temperatures in this study, which may be due to the fish being of similar age and reared under uniform hatchery conditions (Farrell et al., 2009a; Killen et al., 2016). MMR reflects the maximum rate at which aerobic physiological processes can occur, and at 16 °C the salmon may already have been operating near their upper capacity for oxygen uptake and delivery (Clark et al., 2011; McKenzie & Claireaux, 2010; Norin & Clark, 2016; Sandblom et al., 2016). Although no treatment-level differences in MMR were detected, individual variation in thermal reaction norms of SMR and MMR may still exist but could not be resolved with our experimental design. This limitation may help explain why AS did not decline despite SMR increasing while MMR remained stable (Norin et al., 2014; Norin & Metcalfe, 2019; Réveillon et al., 2019). Sample size was reduced from 128 to 72 fish due to lost data, largely caused by malfunction of the water circulation and temperature control system. This uneven sample distribution across temperature treatments may have limited our ability to detect patterns, particularly at 19 and 22 °C. In addition, one extreme outlier was removed from SMR, MMR, and AS analyses based on visual inspection of histograms, and while this was a single fish, its exclusion should be acknowledged. Mortality also occurred in the respirometers (one, three, and two fish at 16, 19, and 22 °C, respectively), and while this represented <10% of the initial sample size for each group, removing these data introduces some potential for survival bias. Importantly, our experiment tested only a single hatchery stock and one juvenile age class; responses could differ in other populations or life stages, especially given that juvenile parr are often more sensitive than eggs or spawners. Together, these limitations highlight the need for caution in extrapolating our findings and suggest that broader testing across stocks, life stages, and larger sample sizes will be important for future studies.

Recent advances in conservation physiology demonstrate how individual-level physiological metrics can be used to guide policy and management strategies, as shown in studies on Sockeye salmon in Canada (Patterson et al., 2016). Integrating TPCs and stock specific information, such as life history genotypes and developmental stages, can inform stakeholders of specific vulnerabilities in populations and how they vary in location and season (Ward et al., 2017). With detailed TPCs, thermal safety margins can be defined in terms of functional success—such as swimming, feeding, and reproducing—rather than mortality alone (Pinsky et al., 2019). Temperature-dependent fisheries management strategies have already been successfully implemented, for example, in Canadian rivers, and could be adapted globally to account for population-specific TPCs (Van Leeuwen et al., 2023). Management strategies, such as adjusting harvest timing and identifying thermal refuges for protection and restoration efforts, could help mitigate the impact of rising temperatures on vulnerable salmon populations.

In recent years, extreme heat events have had catastrophic impacts on migration success and survival across species of salmonids, indicating that a better understanding of their thermal performance is critical for conservation (Baisez et al., 2011; Hinch et al., 2012; Martins et al., 2011, 2012; Muñoz et al., 2021). Our study showed that aerobic scope was not increased between 16 and 22°C in juvenile Atlantic salmon despite a significant increase in SMR. This, combined with the increasing metabolic costs of activities, such as swimming and digestion, indicate that the stable aerobic scope may be leaving juvenile salmon vulnerable to deteriorating performance at temperatures commonly experienced in present-day heatwaves.

## Ethical Declarations

The experiment was performed with permit granted by the Finnish Project Authorisation Board no. ESAVI/16748/2023.

## Acknowledgements

We are grateful for Jonna Hänninen, Juha Hänninen, and staff at Luke Taivalkoski and Luke Laukaa for access to fish and help during the experiment, and for Kari Jauhiainen for technical support, and Prof. Craig Primmer, Dr. Tutku Aykanat, and Dr. Silva Uusi-Heikkilä for access to equipment. We thank Lilian Redon for feedback on an early version of the manuscript.

## Author Contributions

LC and JP: experimental design, conducting experiment, data analysis and writing the manuscript. JP: supervision, management, and funding acquisition. SM: equipment and experimental design. MR and ES: fish care, equipment maintenance and assisting with data collection.

## Data Availability

Data is hosted on Zenodo data repository (https://doi.org/10.5281/zenodo.15308028) and code for analysis is hosted on Github (https://github.com/jprokkola/Salmon-aerobic-scope-2023-public.git).

