## Supplemental material for "Aerobic scope is sustained through a heatwave in juvenile Atlantic salmon"

**Supplementary Figures and Tables**

Lucy Cotgrove^1^, Sergey Morozov^2^, Miika Raitakivi^3^, Evan Sala^3^, Jenni M. Prokkola^1^

**Table S1. Checklist of 53 essential criteria for the reporting of methods for aquatic intermittent-flow respirometry (Killen et al., 2021).**

| Number | **Criterion and Category** | **Response** | **Value (where required)** | **Units** |
| --- | --- | --- | --- | --- |
|  | **EQUIPMENT, MATERIALS, AND SETUP** |  |  |  |
| 1 | Body mass of animals at time of respirometry | Measured Individually after SMR | $\bar{x}$ =1.72 | g |
| 2 | Volume of empty respirometers | Measure by weighing water | $\bar{x}$ =131.86±1 | ml |
| 3 | How chamber mixing was achieved | Circulation loop flowing throughout measurements, discs distributing water in laminar flow |  |  |
| 4 | Ratio of net respirometer volume (plus any associated tubing in mixing circuit) to animal body mass |  | 76:1 |  |
| 5 | Material of tubing used in any mixing circuit | Tygon |  |  |
| 6 | Volume of tubing in any mixing circuit |  | ~12 | ml |
| 7 | Confirm volume of tubing in any mixing circuit was included in calculations of oxygen uptake | Yes |  |  |
| 8 | Material of respirometer (e.g. glass, acrylic, etc.) | Glass |  |  |
| 9 | Type of oxygen probe and data recording | FireStingO2 4-channel oxygen meter and associated optical DO probes and software (Pyro Science GmbH, Aachen, Germany) |  |  |
| 10 | Sampling frequency of water dissolved oxygen | 2 seconds | 2 | second |
| 11 | Describe placement of oxygen probe (in mixing circuit or directly in chamber) | In mixing circuit |  |  |
| 12 | Flow rate during flushing and recirculation, or confirm that chamber returned to normoxia during flushing | Chamber returned to normoxia |  |  |
| 13 | Timing of flush/closed cycles | 5 min open, 15 min closed (16, 19, 22 C) |  |  |
| 14 | Wait (delay) time excluded from closed measurement cycles | Yes | 2 | minutes |
| 15 | Frequency and method of probe calibration (for both 0 and 100% calibrations) | At every temperature | 1 | week |
| 16 | State whether software temperature compensation was used during recording of water oxygen concentration | Yes |  |  |
|  | **MEASUREMENT CONDITIONS** |  |  |  |
| 17 | Temperature during respirometry | Varies according to treatment | 16, 19, 22 | C |
| 18 | How temperature was controlled | Heater/chiller using heated reservoir with temperature controller | ± 0.1C |  |
| 19 | Photoperiod during respirometry | No cover for MMR, kept in dark for SMR |  |  |
| 20 | If (and how) ambient water bath was cleaned and aerated during measurement of oxygen uptake (e.g. filtration, periodic or continuous water changes) | Aerated with air stone and pumped through UV filter, also replaced continuously |  |  |
| 21 | Total volume of ambient water bath and any associated reservoirs | 200 L bath + reservoir | 250 | L |
| 22 | Minimum water oxygen dissolved oxygen reached during closed phases |  | 6.70 mg/L or 70% saturation |  |
| 23 | State whether chambers were visually shielded from external disturbance | Opaque plastic cover for SMR |  |  |
| 24 | How many animals were measured during a given respirometry trial (i.e. how many animals were in the same water bath) | Maximum of 16 per respirometry trial |  |  |
| 25 | If multiple animals were measured simultaneously, state whether they were able to see each other during measurements | No |  |  |
| 26 | Duration of animal fasting before placement in respirometer |  | 24 | hours |
| 27 | Duration of all trials combined (number of days to measure all animals in the study) |  | 22 | days |
| 28 | Acclimation time to the laboratory (or time since capture for field studies) before respirometry measurements |  | 138 | days |
|  | **BACKGROUND RESPIRATION** |  |  |  |
| 29 | State whether background microbial respiration was measured and accounted for, and if so, method used (e.g. parallel measures with empty respirometry chamber, measurements before and after for all chambers while empty, both) | Empty respirometer measurement taken before and after trial |  |  |
| 30 | State if background respiration was measured at beginning and/or end, state how many slopes and for what duration | Beginning and end, 3 slopes |  |  |
| 31 | State how changes in background respiration were modelled over time (e.g. linear, exponential, parallel measures) | Linear |  |  |
| 32 | Level of background respiration (e.g. as a percentage of SMR) | Specifics in online data repository  https://doi.org/10.5281/zenodo.15308028 | $\bar{x}$ = 13.17% |  |
| 33 | Method and frequency of system cleaning (e.g. system bleached between each trial, UV lamp) | System bleached between each temperature change |  |  |
|  | **STANDARD OR ROUTINE METABOLIC RATE** |  |  |  |
| 34 | Acclimation time after transfer to chamber, or alternatively, time to reach beginning of metabolic rate measurements after introduction to chamber | Measurements taken from 4pm onwards |  |  |
| 35 | Time period, within a trial, over which oxygen uptake was measured (e.g. number of hours) |  | ~20 hours |  |
| 36 | Value taken as SMR/RMR (e.g. quantile, mean of lowest 10 percent, mean of all values) | Mean of lowest normal distribution (Chabot 2016) |  |  |
| 37 | Total number of slopes measured and used to derive metabolic rate (e.g. how much data were used to calculate quantiles) | Specifics for each fish in supplementary data files | 31 – 47 total slopes, min. 10% used in SMR calculation. |  |
| 38 | Whether any time periods were removed from calculations of SMR/RMR (e.g. data during acclimation, periods of high activity [e.g. daytime]) | Before 4pm, post MMR |  |  |
| 39 | r^2^ threshold for slopes used for SMR/RMR (or mean) |  | 0.95 |  |
| 40 | Proportion of data removed due to being outliers below r-squared threshold |  | 12 fish (10%) |  |
|  | **MAXIMUM METABOLIC RATE** |  |  |  |
| 41 | When MMR was measured in relation to SMR (i.e. before or after) | Before |  |  |
| 42 | Method used (e.g. critical swimming speed respirometry, swim to exhaustion in swim tunnel, or chase to exhaustion) | Chase to exhaustion |  |  |
| 43 | Value taken as MMR (e.g. the highest rate of oxygen uptake value after transfer, average of highest values) | Immediate **Ṁ**O2 value after transfer |  |  |
| 44 | If MMR measured post-exhaustion, length of activity challenge or chase (e.g. 2 min, until exhaustion, etc.) | 2 minutes |  |  |
| 45 | If MMR measured post-exhaustion, state whether further air-exposure was added after exercise | No further air exposure |  |  |
| 46 | If MMR measured post-exhaustion, time until transfer to chamber after exhaustion or time to start of oxygen uptake recording | Less than 5 seconds |  |  |
| 47 | Duration of slopes used to calculate MMR (e.g. 1 min, 5 min, etc.) | 10 minutes |  |  |
| 48 | Slope estimation method for MMR (e.g. rolling regression, sequential discrete time frames) | spline MMR (Prokkola et al. 2022) |  |  |
| 49 | How absolute aerobic scope and/or factorial aerobic scope is calculated (i.e. using raw SMR and MMR, allometrically mass-adjusted SMR and MMR, or allometrically mass-adjusting aerobic scope itself) | Raw MMR – Raw SMR, then mass adjusted |  |  |
|  | **DATA HANDLING AND STATISTICS** |  |  |  |
| 50 | Sample size |  | 72 |  |
| 51 | How oxygen uptake rates were calculated (software or script, equation, units, etc.) | Script (FishResp, Morozov, 2024) |  |  |
| 52 | Confirm that volume (mass) of animal was subtracted from respirometer volume when calculating oxygen uptake rates | Yes |  |  |
| 53 | State whether analyses accounted for variation in body mass and describe any allometric mass-corrections or adjustments | Yes |  |  |

**Table S2: Number of juvenile *S. salar* included and excluded at each stage of the analysis, divided into temperature groups**.

|  | ***N*** | | | | |
| --- | --- | --- | --- | --- | --- |
| ***Temperature (°C)*** | ***Respirometry experiment*** | ***Dead in chambers*** | ***Lost data*** | ***Excluded from analysis*** | ***Included in statistical analysis*** |
| *16 (week 1)* | 24 | 1 | 0 | 1 | 22 |
| *19* | 32 | 3 | 0 | 10 | 19 |
| *22* | 32 | 2 | 14 | 0 | 16 |
| *16 (week 4)* | 32 | 0 | 16 | 1 | 15 |
| *Total* | 120 | 6 | 30 | 12 | 72 |

**Table S3: Descriptive statistics for absolute metabolic rate measurements (mg O_2_ h^−1^)**. n = number of individuals, x̄ = mean metabolic rate, σ = standard deviation, Min = minimum value, Max = maximum value.

|  |  | **SMR** | | | | **MMR** | | | | **AS** | | | |
| --- | --- | --- | --- | --- | --- | --- | --- | --- | --- | --- | --- | --- | --- |
| **Temperature (°C)** | **n** | $\bar{x}$ | **σ** | **Min** | **Max** | $\bar{x}$ | **σ** | **Min** | **Max** | $\bar{x}$ | **σ** | **Min** | **Max** |
| 16 (week 1) | 22 | 0.27 | 0.08 | 0.17 | 0.47 | 1.16 | 0.56 | 0.22 | 2.59 | 0.89 | 0.50 | 0.03 | 2.14 |
| 19 | 19 | 0.36 | 0.10 | 0.19 | 0.59 | 0.98 | 0.31 | 0.48 | 1.49 | 0.62 | 0.29 | 0.15 | 1.14 |
| 22 | 16 | 0.54 | 0.14 | 0.22 | 0.77 | 1.38 | 0.41 | 0.80 | 2.06 | 0.84 | 0.36 | 0.21 | 1.42 |
| 16 (week 4) | 15 | 0.48 | 0.15 | 0.29 | 0.76 | 1.74 | 0.83 | 0.47 | 3.27 | 1.26 | 0.71 | 0.18 | 2.69 |
| Total | 72 | 0.40 | 0.16 | 0.17 | 0.77 | 1.28 | 0.60 | 0.22 | 3.27 | 0.89 | 0.52 | 0.03 | 2.69 |

**Table S4: Linear equation values for mass adjustment of standard metabolic rate (SMR), maximum metabolic rate (MMR) and aerobic scope (AS; mg O_2_ h^−1^) for equation structure MR = log(weight) + Intercept.** CI indicates 95% confidence intervals for estimates and for model R^2^ for goodness of fit of linear model.

| ***MR*** | ***Intercept*** | ***Lower***  ***Intercept CI*** | ***Upper Intercept CI*** | ***logWeight Estimate*** | ***Lower logWeight CI*** | ***Upper logWeight CI*** | ***R^2^*** |
| --- | --- | --- | --- | --- | --- | --- | --- |
| SMR | -0.612 | -0.654 | -0.570 | 0.859 | 0.698 | 1.021 | 0.616 |
| MMR | -0.141 | -0.198 | -0.084 | 0.978 | 0.757 | 1.199 | 0.526 |
| AS | -0.370 | -0.475 | -0.255 | 1.071 | 0.647 | 1.496 | 0.266 |

**Table S5: Pearson’s correlation coefficient for all metabolic rate measures for juvenile *S. salar* (standard metabolic rate; SMR, maximum metabolic rate: MMR, aerobic scope: AS) at different temperatures.** Table shows both absolute values and mass adjusted values, *r* indicates correlation coefficient and *p* indicates significance. Bolded rows indicate significant relationships with ^*^ *p* < 0.05; ^**^ *p* < 0.01; ^***^ *p* < 0.001.

|  | ***Temperature (°C)*** | ***Variable 1*** | ***Variable 2*** | ***r*** | ***p*** |
| --- | --- | --- | --- | --- | --- |
| Absolute  Values | 16 | **SMR** | **MMR** | **0.8** | **<0.001***** |
|  |  | **SMR** | **AS** | **0.7** | **<0.001***** |
|  |  | **MMR** | **AS** | **0.99** | **<0.001***** |
|  |  | **SMR** | **mass** | **0.95** | **<0.001***** |
|  |  | **MMR** | **mass** | **0.81** | **<0.001***** |
|  |  | **AS** | **mass** | **0.72** | **<0.001***** |
|  | 19 | SMR | MMR | 0.36 | 0.125 |
|  |  | SMR | AS | 0.05 | 0.842 |
|  |  | **MMR** | **AS** | **0.95** | **<0.001***** |
|  |  | **SMR** | **mass** | **0.87** | **<0.001***** |
|  |  | **MMR** | **mass** | **0.65** | **0.002**** |
|  |  | AS | mass | 0.4 | 0.086 |
|  | 22 | **SMR** | **MMR** | **0.51** | **0.042*** |
|  |  | SMR | AS | 0.19 | 0.492 |
|  |  | **MMR** | **AS** | **0.94** | **<0.001***** |
|  |  | **SMR** | **mass** | **0.84** | **<0.001***** |
|  |  | **MMR** | **mass** | **0.78** | **<0.001***** |
|  |  | **AS** | **mass** | **0.55** | **0.026*** |
| Mass Adjusted Values | 16 | SMR | MMR | 0.21 | 0.204 |
|  |  | SMR | AS | 0.15 | 0.370 |
|  |  | **MMR** | **AS** | **0.98** | **<0.001***** |
|  | 19 | SMR | MMR | -0.42 | 0.074 |
|  |  | **SMR** | **AS** | **-0.53** | **0.018*** |
|  |  | **MMR** | **AS** | **0.98** | **<0.001***** |
|  | 22 | SMR | MMR | -0.37 | 0.159 |
|  |  | **SMR** | **AS** | **-0.58** | **0.020*** |
|  |  | **MMR** | **AS** | **0.95** | **<0.001***** |

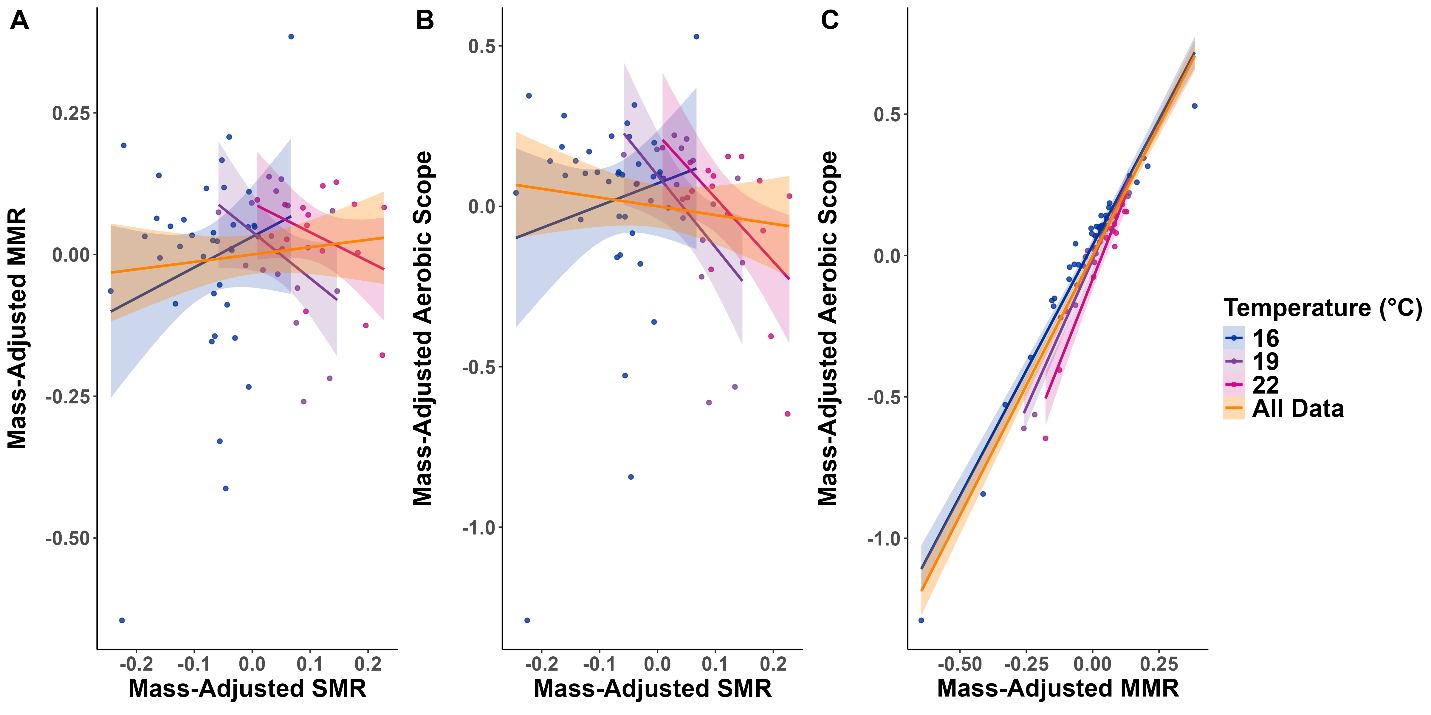

**Figure S1: Scatter plots showing the relationship of mass adjusted metabolic rates for juvenile *S. salar*.** **A**: Standard metabolic rate (SMR) and maximum metabolic rate (MMR), B: SMR and aerobic scope, **C**: MMR and aerobic scope. Each point represents individual fish, with different colours representing different tested temperatures (16 °C: Blue, 19 °C: Purple, 22 °C: Pink). Goodness of fit lines indicate linear regression and shading shows 95% confidence intervals for each temperature, with orange colour representing fit of all data.
